# High-grade serous ovarian cancer development and anti-PD-1 resistance is driven by IRE1α activity in neutrophils

**DOI:** 10.1101/2024.08.05.606646

**Authors:** Alexander Emmanuelli, Camilla Salvagno, Sung Min-Hwang, Deepika Awasthi, Tito A. Sandoval, Chang-Suk Chae, Jin-Gyu Cheong, Chen Tan, Takao Iwawaki, Juan R. Cubillos-Ruiz

## Abstract

High-grade serous ovarian cancer (HGSOC) is an aggressive malignancy that remains refractory to current immunotherapies. While advanced stage disease has been extensively studied, the cellular and molecular mechanisms that promote early immune escape in HGSOC remain largely unexplored. Here we report that primary HGSO tumors program neutrophils to inhibit T cell anti-tumor function by activating the endoplasmic reticulum (ER) stress sensor IRE1α. We found that intratumoral neutrophils exhibited overactivation of ER stress response markers compared with their counterparts at non-tumor sites. Selective deletion of IRE1α in neutrophils delayed primary ovarian tumor growth and extended the survival of mice with HGSOC by enabling early T cell-mediated tumor control. Notably, loss of IRE1α in neutrophils sensitized tumor-bearing mice to PD-1 blockade, inducing HGSOC regression and long-term survival in ∼50% of treated hosts. Hence, neutrophil-intrinsic IRE1α facilitates early adaptive immune escape in HGSOC and targeting this ER stress sensor might be used to unleash endogenous and immunotherapy-elicited immunity that controls metastatic disease.

## Introduction

High-grade serous ovarian carcinoma (HGSOC) is a common sub-type of epithelial ovarian cancer and a leading cause of lethal gynecologic malignancy in the United States (Momenimovahed et al., 2019; Siegel et al., 2023). Due to chemotherapy resistance, high recurrence rates, and few alternative treatment options, there is an urgent need to develop novel approaches to tackle this disease (Matulonis et al., 2016). While recent advances in targeted therapy using poly adenosine diphosphate-ribose polymerase (PARP) inhibitors show potential, the implementation of immune checkpoint blockade has been far less successful, despite the positive association of T cell infiltration with survival in HGSOC patients (Hwang et al., 2012; Konstantinopoulos et al., 2020; Matulonis et al., 2019). This clinical challenge indicates that alternative mechanisms of T cell inhibition remain actively engaged in the environment of HGSOC. Identifying, understanding, and targeting these mechanisms is crucial to improve the effectiveness of T cell-based immunotherapy in women with HGSOC.

Neutrophils are an abundant innate immune cell, long known for their role as a first line of defense against pathogens (Jones et al., 2016; Kolaczkowska and Kubes, 2013; Rosales, 2018). Recently, neutrophils have emerged as critical mediators of immune regulation and inflammation in the tumor microenvironment (TME), and as key players in cancer initiation, progression, and metastasis to distant anatomical sites (Coffelt et al., 2016; Jaillon et al., 2020; Mayer et al., 2016). Several studies have described tumor-associated neutrophils (TANs) as a highly immunosuppressive and pro-tumorigenic subset of myeloid cells that stifle adaptive T cell responses (Condamine et al., 2016; Kim et al., 2022; Siolas et al., 2021; Wang et al., 2023). Despite strong evidence of their pro-tumorigenic nature, studies of TANs in a variety of cancer models have showcased their malleability in this role. For instance, in mouse models of melanoma and lung cancer, tumor growth is drastically hindered by the priming of anti-tumor neutrophil responses via β-glucan-induced trained granulopoiesis (Kalafati et al., 2020). A seminal study of human non-small cell lung cancer demonstrated that patient-derived neutrophils have the capacity to stimulate T cell activity at early stages of disease (Eruslanov et al., 2014). Human polymorphonuclear neutrophils can release cancer cell killing factors and reduce tumor burden in patient-derived xenograft models of breast and lung cancers, though no analog has been identified in mice (Cui et al., 2021). Neutrophil infiltration in human ovarian cancers correlate with poor patient prognosis, while in an transplantable model of murine ovarian cancer, production of neutrophil extracellular traps (NETs) was critical in the formation of the pre-metastatic tumor niche (Jeerakornpassawat and Suprasert, 2020; Lee et al., 2019; Patel et al., 2018). The diverse roles that neutrophils play in the progression of cancer highlights their plasticity, yet the signaling pathways by which TANs sense and interpret the stimuli they encounter in the TME are still insufficiently understood.

Dysregulated endoplasmic reticulum (ER) stress responses have emerged as central molecular regulators of immune cell function in the TME (Chen and Cubillos-Ruiz, 2021; Cubillos-Ruiz et al., 2017; Di Conza et al., 2023; Salvagno et al., 2022). ER stress triggers the unfolded protein response (UPR), an adaptive pathway that operates to restore ER homeostasis. The UPR is initiated and governed by three ER stress sensors: Inositol-requiring enzyme 1 α (IRE1α), protein kinase R-like endoplasmic reticulum kinase (PERK), and activating transcription factor 6 (ATF6) (Hetz et al., 2020). IRE1α is a serine/threonine kinase that upon detection of accumulated misfolded proteins undergoes dimerization, auto-phosphorylation, and activation of its endoribonuclease domain (Hetz et al., 2020). These conformational changes allow IRE1α to cleave the primary X-box binding protein (*XBP1)* mRNA transcript in the cytosol, converting it into a spliced isoform (*XBP1s*) that is translated into the functionally active transcription factor, XBP1s (Yoshida et al., 2001). Canonically, XBP1s upregulates a variety of chaperones and foldases to restore ER proteostasis (Lee et al., 2003). However, it is now well appreciated that IRE1α-XBP1s signaling can modulate UPR-independent transcriptional and metabolic pathways in a cell-specific and context-dependent manner. Consequently, UPR activation can govern cellular phenotypes and fates implicated in tumor development and metastasis (Crowley et al., 2023; Cubillos-Ruiz et al., 2015; Gaudette et al., 2020; Logue et al., 2018; Song and Cubillos-Ruiz, 2019). The expression of UPR-associated genes has been correlated with immunosuppressive polymorphonuclear myeloid-derived suppressor cells in non-small cell lung cancer as well as head and neck cancer patients (Condamine et al., 2016), but whether ER stress pathways endow neutrophils with pro-tumoral phenotypes is unknown.

Here we report that IRE1α activation drives immunosuppressive reprogramming of TANs in HGSOC, which promotes disease progression by inhibiting T cell-dependent anti-tumor immunity. Ablating IRE1α in neutrophils unleashes long-term T cell-mediated control of HGSOC and sensitizes hosts with this disease to immune checkpoint blockade.

## Results

### Neutrophils promote primary tumor growth in an autochthonous mouse model of HGSOC

Mounting evidence supports the hypothesis that HGSOC precursor lesions originate from epithelial cells in the fallopian tube, which eventually spread to the ovary leading to the formation of solid tumors (Kurman and Shih Ie, 2016; Piek et al., 2001). While extensive research of metastatic epithelial ovarian cancer has been carried out with transplantable syngeneic cell lines (e.g. the ID8 model), these fail to recapitulate the earliest events of tumorigenesis that dictate the development of the disease. Although neutrophils and macrophages have been implicated in transcoelomic spread, it is unclear what role they play in HGSOC tumor initiation (Lee et al., 2019; Yin et al., 2016). Thus, we utilized a recently developed autochthonous model to induce HGSOC in female mice by direct injection of non-viral plasmid constructs enabling CRISPR/Cas9-mediated loss of p53 and concomitant transgenic overexpression of c-Myc into the ovary and fallopian tube, followed by electroporation (**Fig 1A**) (Paffenholz et al., 2022). This microsurgical approach recapitulates the co-occurrence of *MYC* amplifications and *TP53* mutations found in ∼35% of patients with HGSOC, and results in the development of primary ovarian tumors with subsequent ascites formation, peritoneal carcinomatosis, and omental metastasis (Cancer Genome Atlas Research, 2011; Lengyel, 2010; Vang et al., 2016).

**Figure 1:**
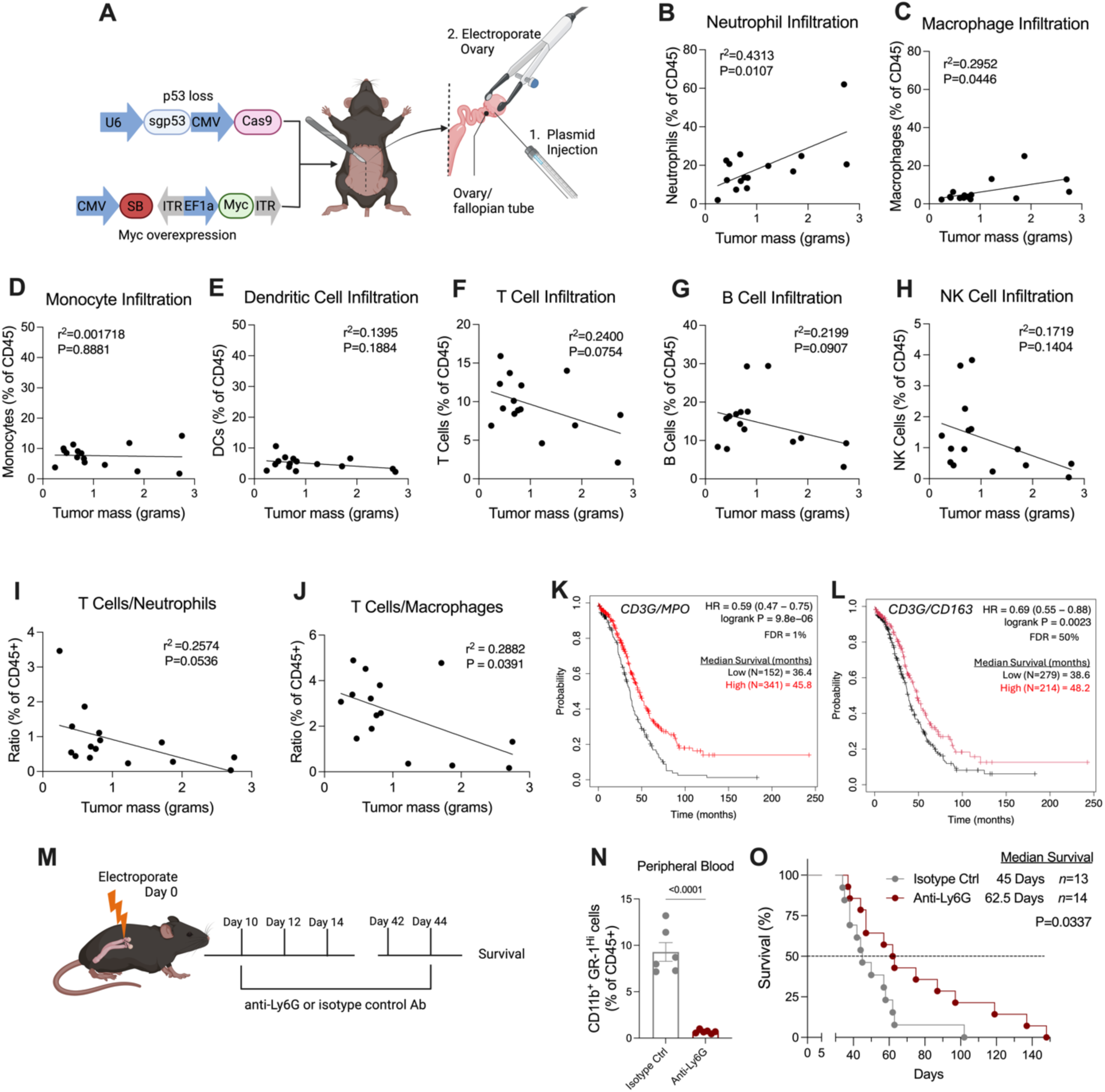
Neutrophils facilitate tumor progression in an autochthonous model of HGSOC. (**A**) Schematic of the experimental system to generate autochthonous HGSOC. Plasmid vectors enabling Myc overexpression and CRISPR-Cas9-mediated targeting of *Trp53* are co-delivered into the ovary by *in vivo* electroporation. (**B-H**) Correlation plot between the mass of the primary ovarian tumor of C57BL/6J mice and infiltrating immune cell populations based on FACS analysis. (**B**) Neutrophils (CD45^+^CD11b^+^Ly6G^Hi^Ly6C^Lo^), (**C**) Macrophages (CD45^+^CD11b^+^F4/80^+^), (**D**) Monocytes (CD45^+^CD11b^+^Ly6G^-^ Ly6C^+^), (**E**) Dendritic cells (CD45^+^CD11c^+^MHC-II^+^), (**F**) T cells (CD45^+^CD3^+^), (**G**) B cells (CD45^+^CD19^+^), (**H**) NK cells (CD45^+^NK1.1^+^). Correlation plots between mass of the primary tumor and the proportional ratio of (**I**) T cell to neutrophils and (**J**) T cells to macrophages. Human HGSOC patient survival analysis based on bulk RNA-seq TCGA data of the indicated genes. Patients are assessed based on relative ratio of (**K**) intratumoral T cells (*CD3G*) to neutrophils (*MPO*) and (**L**) intratumoral T cells (*CD3G*) to macrophages (*CD163*). Patients with low ratio of gene expression are compared to those with a high ratio score. (**M**) I*n vivo* antibody-mediated depletion scheme. 10 Days after autochthonous HGSOC inductionC57BL/6J mice were electroporated with oncogenic plasmids. At Day 10, mice were treated as indicated in the figure. (**N**) Flow cytometry quantification of neutrophils in peripheral blood. (**O**) Overall survival of ovarian tumor-bearing C57BL/6J mice that were i.p. injected with neutrophil-depleting antibody (anti-Ly6G) or an isotype control (IgG2a). (**B-J**) Simple linear regression, Pearson correlation coefficient (r^2^) and P-value provided (*n* =15 mice). (**K-L**) Cox proportional hazards model, results were adjusted for false discovery rate (FDR). (**N**) Unpaired t-test (*n* =6). (**O**) Log-Rank (Mantel Cox) test (IgG2a *n*= 13, αLy6G *n* =14) exact P-values are provided.

Unique cancer types produce distinct TMEs with diverse immune infiltrates and cellular states (Binnewies et al., 2018). To gain insight into the cellular composition of the primary ovarian tumors generated in the autochthonous HGSOC system, we resected tumors at various disease stages and analyzed immune cell infiltration by flow cytometry (**Fig S1A**). We found that as primary tumor size increased, the proportion of CD45^+^ infiltrates markedly skewed towards neutrophils (**Fig 1B**). The proportion of macrophages also increased with the primary tumor mass, while monocyte and dendritic cell infiltration remained relatively constant (**Fig 1C-E**). Of note, the proportion of lymphocytes, particularly T cells, tended to decrease (**Fig 1F-H**). Further, we observed that neutrophils were the dominant myeloid subset in hemorrhagic ascites, but not in omental metastatic lesions of HGSOC bearing mice (**Fig S1B-C**). A high neutrophil-to-lymphocyte ratio is a strong, independent predictive biomarker of clinical outcome in ovarian cancer patients (Cho et al., 2009; Liontos et al., 2021; Williams et al., 2014). Similarly, we identified a negative association between T cell infiltration and the proportion of both neutrophils and macrophages in autochthonous ovarian tumors (**Fig 1I-J**). Furthermore, by analyzing bulk RNA-seq data from 493 HGSOC patients in The Cancer Genome Atlas (TCGA), we found that high expression ratios of *CD3G* (indicative of T cell infiltration) to myeloperoxidase (*MPO*) transcripts, mainly expressed by neutrophils, significantly correlated with increased overall survival rates (**Fig 1K**) (Cancer Genome Atlas Research, 2011; Gyorffy, 2023). However, there was a weak correlation between high ratios of *CD3G* to *CD163*, mainly expressed by macrophages, with increased overall survival (**Fig 1L**). Robust neutrophil recruitment correlating with tumor size, decreased T cell infiltration, and disease progression prompted us to speculate that these myeloid cells play a critical role in the autochthonous model of HGSOC.

Lee *et al* reported that neutrophil depletion in mice hinders tumor progression in a genetically distinct, orthotopic model of ovarian cancer (Lee et al., 2019). Moreover, Siolas *et al* demonstrated that the function of TANs in cancer models is drastically impacted by the tumor genotype (Siolas et al., 2021). Thus, to determine the role of neutrophils in autochthonous HGSOC development we employed an antibody-mediated depletion scheme. Ten days after C57BL/6J mice were induced with HGSOC, mice were randomized into two groups that were treated for five weeks with intraperitoneal (i.p.) injections of an anti-Ly6G depleting antibody or its corresponding isotype control antibody (**Fig 1M**). Neutrophil depletion efficacy was confirmed by flow cytometry analysis of peripheral blood (**Fig 1N**). Of note, mice treated with anti-Ly6G had a significant increase in median survival of 16 days compared with their control counterparts (**Fig 1O**). These data reveal that neutrophils are crucial for the optimal initiation and progression of HGSOC in the autochthonous model utilized.

### Tumor-associated neutrophils exhibit overexpression of ER stress gene markers

Tumor-infiltrating immune cells face persistent, harsh conditions that disrupt ER homeostasis and thus activate the UPR (Chen and Cubillos-Ruiz, 2021; Cubillos-Ruiz et al., 2017; Di Conza et al., 2023; Salvagno et al., 2022). Indeed, intratumoral ER stress and UPR activation have emerged as predictors of clinical outcomes in cancer patients and drivers of immunosuppression and disease progression in diverse preclinical models of cancer (Cubillos-Ruiz et al., 2015; Samanta et al., 2020; Song and Cubillos-Ruiz, 2019). Aberrant activation of ER stress sensing pathways has been documented in ovarian tumor-infiltrating T cells (Song et al., 2018) and dendritic cells (Cubillos-Ruiz et al., 2015), but whether neutrophils undergo ER stress responses implicated in HGSOC initiation and progression remains elusive. We sought to evaluate the activation status of the UPR in tumor-associated neutrophils (TANs). To this end, we collected bone marrow, peripheral blood, spleen, primary ovarian tumor, ascitic fluid, and omental metastases from female mice bearing HGSOC. Neutrophils were then FACS sorted from each tissue for expression analysis of ER stress-related gene markers. We found that TANs demonstrated high *Xbp1* mRNA splicing (*Xbp1s/Xbp1^Total^*), indicative of IRE1α activation, and overexpression of canonical UPR gene markers controlled by XBP1s such as *Dnajb9* and *Hspa5,* compared with neutrophils from the bone marrow (**Fig 2A-C**). We also observed increased expression of *Ddit3* in TANs, suggesting activation of the PERK arm of the UPR (**Fig 2D**). Of note, there was a positive correlation between the expression of *Dnajb9* and *Hspa5*, but not *Ddit3*, with high levels of *Xbp1s/Xbp1^total^* (**Fig 2E-G**). These data indicate that neutrophils at the tumor site exhibit IRE1α overactivation.

**Figure 2:**
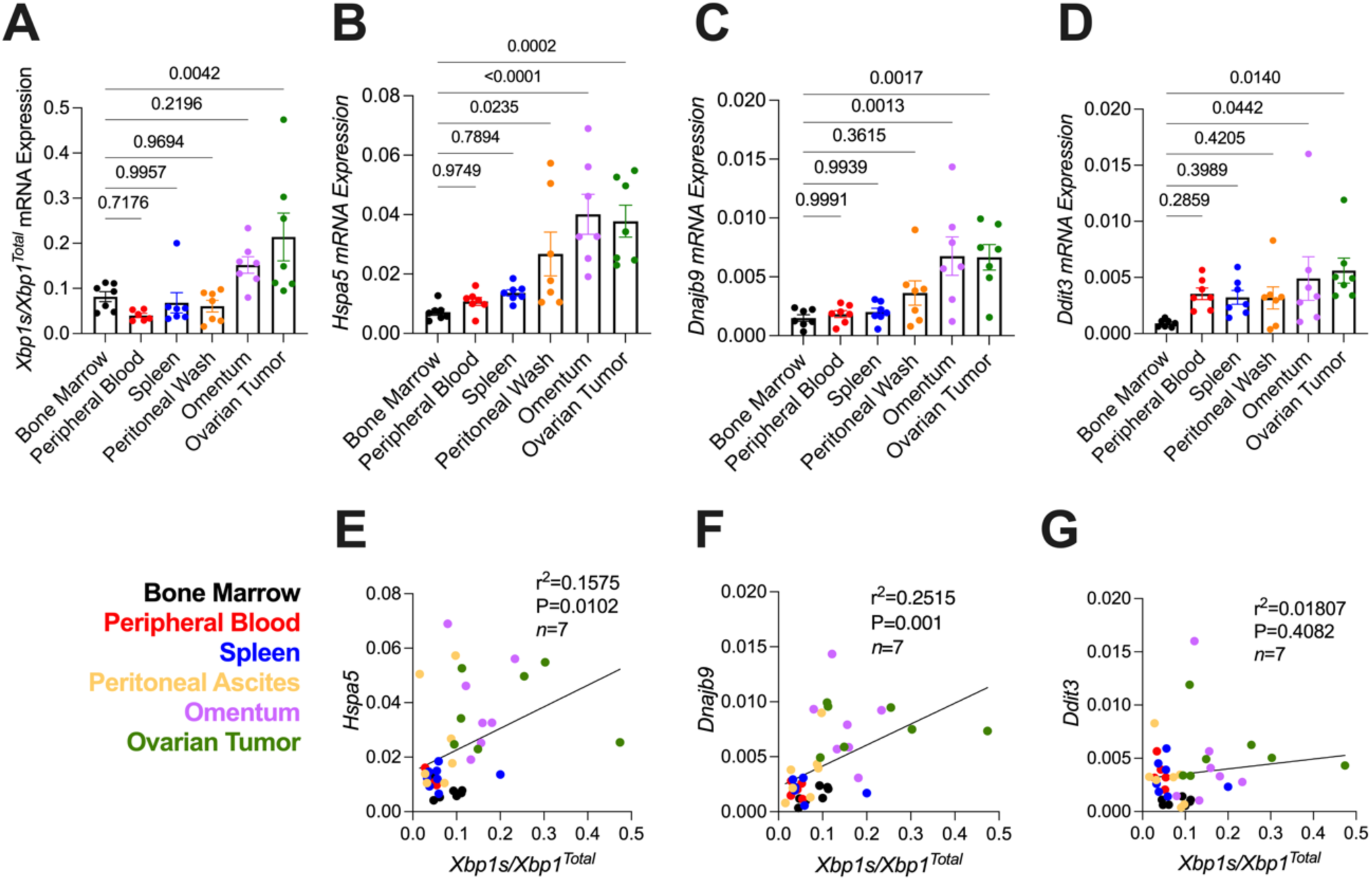
Enhanced IRE1α-XBP1s activation in intratumoral neutrophils. C57BL/6J mice were induced with autochthonous HGSOC as described in figure 1A. (**A-D**) CD45^+^CD11c^-^CD19^-^F4/80^-^ CD11b^+^Ly6G^Hi^Ly6C^Lo^ TANs were FACS sorted from tumor bearing mice and expression of indicated genes was analyzed by RT-qPCR. (**E-G**) Correlation of ER stress gene expression in neutrophils from tumor bearing mice. (**A-D**) One-way ANOVA (*n* =7 mice). (**E-G**) Pearson correlation coefficient (r^2^) and P-value (*n* =7 mice) provided.

### Selective ablation of IRE1**α** in neutrophils delays the progression of HGSOC and extends host survival

We next sought to determine if neutrophil-intrinsic IRE1α was required for the optimal development of autochthonous HGSO tumors. To this end, we crossed *Ern1^f/f^* mice with the *Mrp8^Cre^* strain that enables selective gene deletion in neutrophils (Abram et al., 2014). Autochthonous HGSOC was developed in these conditional knockout mice (*Ern1^f/f^ Mrp8^Cre^*) and their IRE1α-sufficient (*Ern1^f/f^*) counterparts. RT-qPCR analyses confirmed that TANs isolated from *Ern1^f/f^ Mrp8^Cre^* developing HGSOC failed to overexpress *Xbp1s* (**Fig 3A**), indicating effective IRE1α ablation in this population. Further, the *Mrp8^Cre/+^* strain harbors a GFP reporter that is selectively expressed in neutrophils, which we confirmed using total splenocytes and malignant cells from *Ern1^f/f^* and *Ern1^F/F^ Mrp8^Cre^* tumor-bearing mice (**Fig S2A-B**). Flow cytometry analysis of TAN infiltration in the primary ovarian tumor, ascites, and omental metastases from mice in the late stages of disease showed that neutrophil infiltration into these tissues was unaffected by the deletion of IRE1α (**Fig 3B**). Similarly, neutrophil infiltration into the primary ovarian tumor site as a function of tumor growth showed no significant difference between the two groups (**Fig 3C**). Expression of chemokine receptors CCR2 and CXCR3, both previously highlighted as functional markers of TANs, were comparable in neutrophils isolated from *Ern1^f/f^* versus *Ern1^f/f^ Mrp8^Cre^* mice with HGSOC (**Fig S2C-D**) (Bonecchi et al., 2010; Massara et al., 2018). We quantified cell surface expression of other common neutrophil chemokine receptors including CCR1, CXCR4, and CXCR5, but observed few cells that expressed these markers (not shown). These data indicate that IRE1α is dispensable for the survival or normal infiltration of neutrophils into primary ovarian tumors.

**Figure 3:**
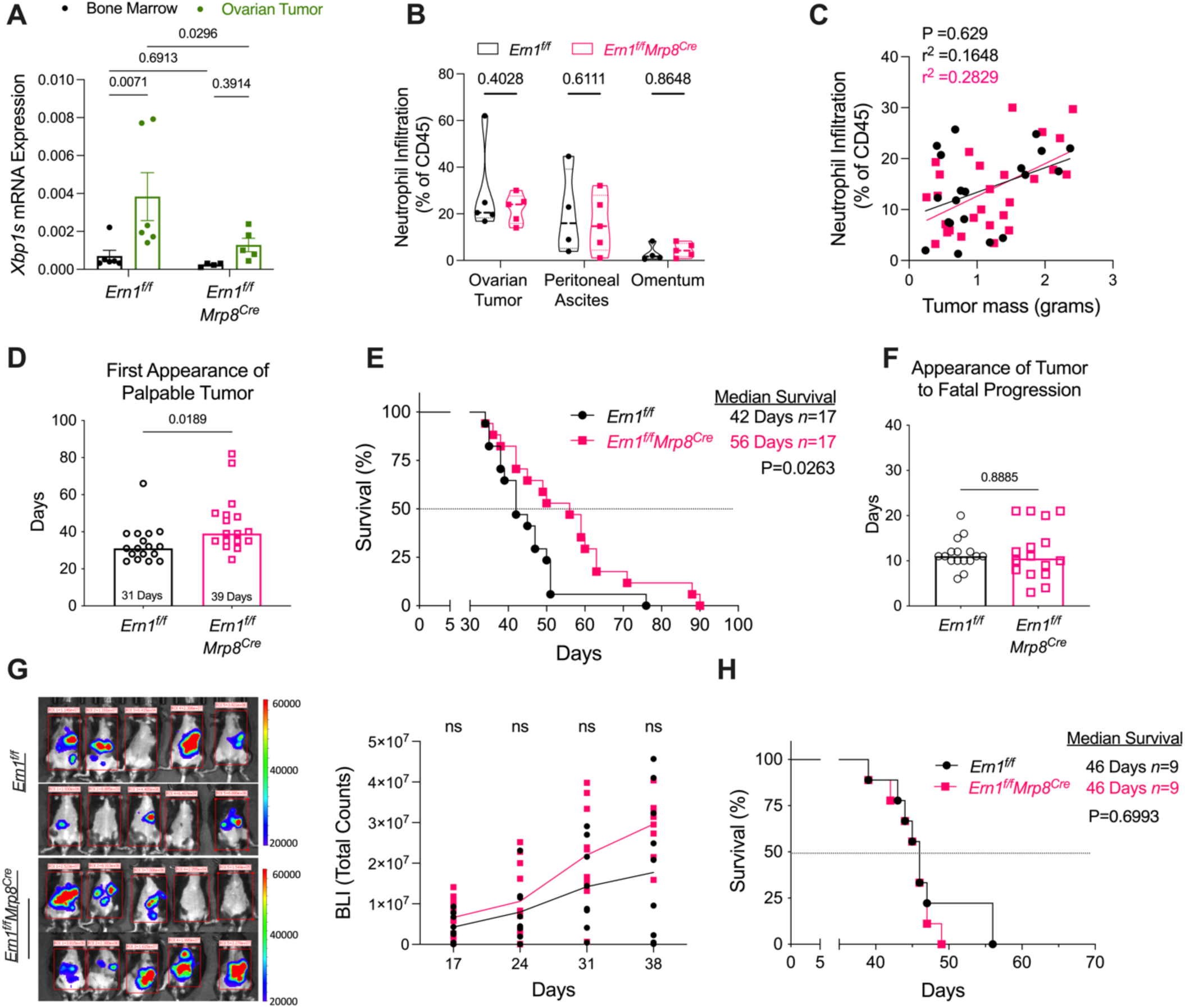
Loss of IRE1α in neutrophils blunts primary HGSOC progression. (**A-F**) Analysis *of Ern1^f/^*^f^ and *Ern1^f/f^* Mrp8^Cre^ mice bearing autochthonous HGSOC. (**A**) RT-qPCR analysis of neutrophils sorted from the bone marrow and primary ovarian tumor of mice with autochthonous HGSOC. (**B**) Violin plots showing the proportion of neutrophils in the primary ovarian tumor, ascites, and omental metastases. (**C**) Correlation between primary ovarian tumor mass and neutrophil infiltration by flow cytometry. (**D-F**) Tumor bearing mice were palpated bi-weekly from day 21 post-surgery to determine (**D**) median time to develop ovarian tumors, (**E**) Kaplan-Meier survival curve of overall survival rates, and (**F**) median time of the progression of HGSOC tumor growth. (**G-H**) *Ern1^f/f^* or *Ern1^f/f^ Mrp8^Cre^* mice implanted i.p. with the transplantable MP cell line. (**G**) Metastatic progression of tumor-bearing mice. *Left:* Representative BLI of tumor-bearing mice at day 24 post-implantation. *Right:* Quantification of peritoneal tumor burden over time. (**H**) Kaplan-Meier survival curve of MP-bearing mice. (**A-B**) 2-way ANOVA (*n* =4-6 mice/group). (**C**) Simple linear regression, Pearson correlation coefficient (r^2^) and P-value provided (*Ern1^f/f^ n* =21, *Ern1^f/f^ Mrp8^Cre^ n* =27). (**D, F**) Unpaired t-test (*n* =17/group). (**G**) 2-way ANOVA (*Ern1^f/f^ n* =10, *Ern1^f/f^ Mrp8^Cre^ n* =10). (**E, H**) Log-Rank (Mantel Cox) test. Group numbers and P*-*values provided.

We next monitored tumor growth and overall survival in *Ern1^f/f^* and *Ern1^f/f^ Mrp8^Cre^* mice developing autochthonous HGSOC. Strikingly, we found that selective IRE1α deletion in neutrophils delayed the onset of HGSOC and therefore extended overall host survival by 14 days (**Fig 3D-E**). While mice lacking IRE1α in neutrophils demonstrated delayed onset of primary tumors (**Fig 3D**), the absence of this sensor did not appear to affect tumor progression or metastasis upon entering advanced disease stages (**Fig 3F**). We performed similar survival experiments utilizing *Ern1^f/f^LysM^Cre^* mice that preferentially delete IRE1α in macrophages (Abram et al., 2014; Batista et al., 2020) but did not observe a significant survival benefit compared to their control littermates (**Fig S3A**). Collectively, these data reveal that neutrophil-intrinsic IRE1α is necessary for the optimal initiation and progression of primary HGSOC.

We then evaluated whether IRE1α expression in neutrophils also facilitated disease progression in transplantable models of advanced ovarian carcinoma. To this end, we used the MP cell line derived from late-stage, metastatic autochthonous HGSOC tumors (Paffenholz et al., 2022). Notably, we found no differences in peritoneal metastatic progression or overall survival in *Ern1^f/f^* versus *Ern1^f/f^ Mrp8^Cre^* mice challenged with this transplantable cell line (**Fig 3G-H**). Similar results were observed in two additional transplantable models of metastatic ovarian cancer: the *Trp53^R172H^Pten^-/-^Nf1^-/-^Myc^OE^* (PPNM) model of high-grade tubo-carcinoma (**Fig S3B**) and the ID8-*Defb29/Vegf-A* system (**Fig S3C-D**) (Conejo-Garcia et al., 2004a; Iyer et al., 2021). These results reveal a salient yet contextual role for neutrophil-intrinsic IRE1α specific to the initiation and early progression of autochthonous HGSOC.

### Ablating IRE1**α** in neutrophils unleashes T cell-mediated control of HGSOC

Myeloid cells can dictate the phenotype of anti-tumor T cells (Conejo-Garcia et al., 2004b; Cubillos-Ruiz et al., 2012; Puttock et al., 2023), but the precise molecular mechanisms by which neutrophils hinder protective lymphocyte populations in primary HGSOC remain obscure. Hence, we next sought to determine if the survival benefit observed in *Ern1^f/f^ Mrp8^Cre^* mice developing HGSOC was dependent on enhanced T cell-mediated anti-tumor responses. We employed antibody-based depletion of CD4 and CD8 T cells in tumor bearing mice. Ten days after autochthonous HGSOC induction, *Ern1^f/f^* and *Ern1^f/f^ Mrp8^Cre^* mice received bi-weekly i.p. injections of both anti-CD4 and anti-CD8 depleting antibodies, or an isotype control IgG2b antibody, until the survival endpoint was reached. Successful depletion was confirmed by analysis of T cell populations in peripheral blood (**Fig 4A**). While the ablation of T cells did not affect the overall survival of *Ern1^f/f^* mice bearing HGSOC, it completely abrogated the beneficial effect conferred in *Ern1^f/f^ Mrp8^Cre^* hosts (**Fig 4B**). These data demonstrate that T cells control primary HGSOC progression when IRE1α expression has been ablated in neutrophils.

**Figure 4:**
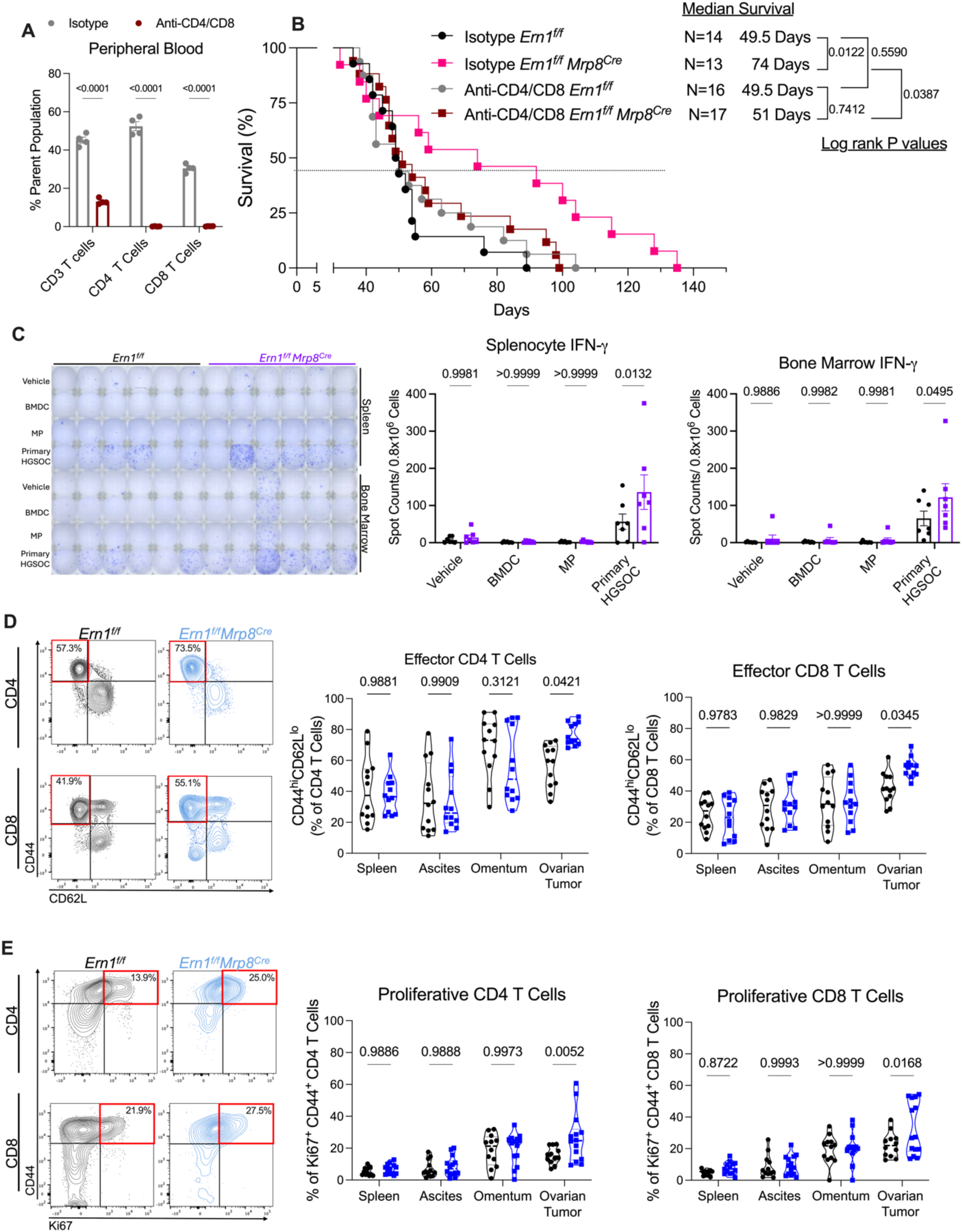
Neutrophil-intrinsic IRE1α curtails T cell-dependent control of primary HGSOC: (**A**) Flow cytometry-based analysis of T cell populations found in peripheral blood 48 hours after i.p. administration of T cell-depleting antibodies (anti-CD4/CD8) or an isotype (IgG2b) control. (**B**) Kaplan-Meier survival curves for the indicated treatment groups. (**C-E**) Indicated tissue samples were harvested from autochthonous HGSOC-bearing mice (**C**) Total splenocytes or bone marrow cells were seeded in a 96-well plate and challenged with the indicated lysates. *Left:* Representative image of IFN-γ ELISpot assay and *right:* Quantification of results. (**D-E**) CD4 and CD8 T cells (CD45^+^CD11b^-^CD11c^-^CD19^-^CD3^+^) analyzed by flow cytometry. (**D**) *left* Representative FACS plots and corresponding violin plots for the proportion of *center:* CD44^Hi^ CD62L^lo^ effector CD4^+^ T cells and *right:* effector CD8^+^ T cells. (**E**) *left* representative FACS plots and corresponding violin plots for the proportion of *center:* CD44+ Ki67+ CD4^+^ and *right:* CD8^+^ T cells. (**A**) 2-way ANOVA (*Ern1^f/f^ n* =4, *Ern1^f/f^ Mrp8^Cre^ n* =4). (**B**) Log-Rank (Mantel Cox) test, P-value and number of mice per group provided. (**C**) 2-way ANOVA (*n* =7/group). (**D**) 2-way ANOVA (*n* =12/group). (**E**) 2-way ANOVA (*Ern1^f/f^ n* =12, *Ern1^f/f^ Mrp8^Cre^ n* =14). P-values provided.

Previous studies of spontaneous T cell reactivity to tumor neoantigens in human and mouse ovarian cancers have reported mixed results where variable mutational burden or the low frequency of reactive T cells limit cancer immunosurveillance (Bobisse et al., 2018; Liu et al., 2019; Martin et al., 2016; Wick et al., 2014). Yet, in our autochthonous HGSOC system, the survival benefit observed in *Ern1^f/f^ Mrp8^Cre^* mice was entirely dependent on CD4 and CD8 T cells, implying a robust tumor antigen-specific response. Lymphoid organs such as bone marrow and spleen can serve as reservoirs of antigen-experienced T cell populations in cancer (Mahnke et al., 2005). Thus, leukocytes from these tissues were isolated from HGSOC-bearing mice and exposed to lysates generated from autologous bone marrow-derived dendritic cells (BMDCs, negative control), the MP ovarian cancer cell line, or primary autochthonous tumors, for subsequent analysis of IFN-γ production in ELISPOT assays. Lymphocytes from the spleen and bone marrow of *Ern1^f/f^ Mrp8^Cre^* mice demonstrated significantly higher reactivity to autochthonous tumor lysates than their wild type counterparts (**Fig 4C**). We observed negligible reactivity in both groups upon stimulation with protein lysates from the MP cell line or control autologous BMDCs, indicating that memory T cells from HGSOC-bearing mice produce IFN-γ only upon recall by their cognate antigens (**Fig 4C**). Hence, neutrophil-specific deletion of IRE1α enables the generation of systemic memory immune responses against primary tumor growth that is evidenced in distal sites such as the bone marrow and spleen.

We next examined the intratumoral T cell activation status in *Ern1^f/f^* or *Ern1^f/f^ Mrp8^Cre^* mice bearing autochthonous HGSOC. We evaluated spleen, ascites, metastasized omental tissue and primary ovarian tumors. While there was a marked increase in the proportion of antigen-experienced effector (CD44^+^ CD62L^-^) CD4 and CD8 T cells in primary ovarian tumors of *Ern1^f/f^ Mrp8^Cre^* hosts, we observed no differences in metastasized tissues or the spleen (**Fig 4D**). To expand these findings, we assessed expression of the proliferation marker Ki67 and found a significant increase in both CD4 and CD8 T cells infiltrating primary ovarian tumors of *Ern1^f/f^ Mrp8^Cre^* mice compared with their control counterparts (**Fig 4E**). No differences were found in other sites (**Fig 4E**), indicating changes that are selective to the primary tumor site.

Increased activation of intratumoral T cells in *Ern1^f/f^Mrp8^Cre^* mice indicated that they possess enhanced effector function. Therefore, we analyzed their capacity to produce IFN-γ and TNF-α in situ. We observed no differences in IFN-γ or TNF-α expression by intratumoral CD4 or CD8 T in mice of either genotype (**Fig S4A-C**). We also analyzed the proportion of CD107a^+^ CD8 T cells but few expressed this cytotoxic degranulation marker (data not shown). The effector status of T cells infiltrating primary tumors in *Ern1^f/f^ Mrp8^Cre^* mice differed from our observation of their enhanced activation and proliferative capacity (**Fig. 4, D-E**). Therefore, we surmised that parallel mechanisms of immune regulation, beyond neutrophil-intrinsic IRE1α, could inhibit intratumoral T cell function and T cell-mediated tumor control, enabling immune escape and fatal disease progression.

### Loss of IRE1**α** in neutrophils sensitizes HGSOC to PD-1 blockade

Ablating IRE1α in neutrophils curtailed ovarian tumorigenesis by activating adaptive anti-tumor immunity. However, even in an immunogenic setting, tumor-infiltrating T cells can enter into a state of exhaustion/dysfunction that allows cancers to progress (Scott et al., 2019). We hypothesized that while deleting IRE1α in neutrophils activates relevant anti-HGSOC immune responses, T cells infiltrating these tumors are affected by additional immunosuppressive mechanisms enriched in the TME, such as engagement of immune checkpoints (Ephraim et al., 2022). Thus, we analyzed the surface expression of PD-1, CTLA4, and Lag-3 on intratumoral T cells from *Ern1^f/f^* and *Ern1^f/f^ Mrp8^Cre^* mice with autochthonous HGSOC. Notably, CD4 and CD8 T cells from the primary tumor drastically upregulated PD-1 and Lag-3, but not CTLA4, compared to splenic T cells, in both *Ern1^f/f^ Mrp8^Cre^* and *Ern1^f/f^* mice (**Fig 5A-B, Fig S5A-B**). Furthermore, 40% of T cells in these primary tumors were PD-1^+^, suggesting they may remain susceptible to inhibitory PD-1/PD-L1 signaling throughout malignant progression despite IRE1α deletion in neutrophils (Dumitru et al., 2022). Indeed, we found that total CD11b^+^ myeloid populations in primary tumors from both *Ern1^f/f^* and *Ern1^f/f^ Mrp8^Cre^* female mice expressed high levels of PD-L1, relative to their counterparts in distal sites (**Fig 5C**). High PD-1 expression by intratumoral T cells and enrichment of PD-L1 in myeloid cells present in the same milieu prompted us to speculate that IRE1α-deletion in neutrophils and concomitant PD-1 blockade could induce synergistic anti-HGSOC effects.

**Figure 5:**
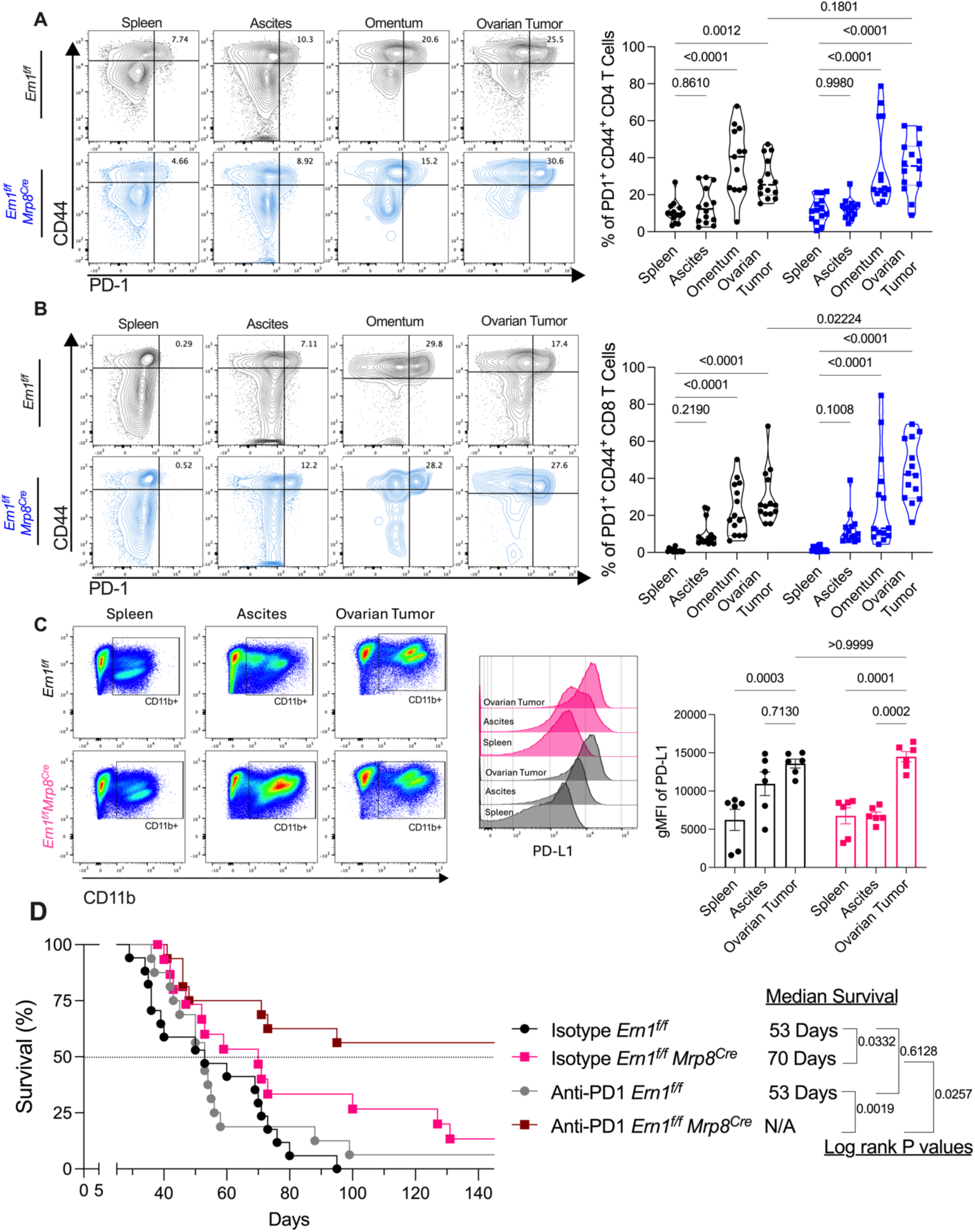
Ablation of IRE1α in neutrophils sensitizes HGSOC hosts to PD-1 blockade immunotherapy: Female mice were induced with autochthonous HGSOC as described in Figure 1A and the indicated tissue samples were harvested from tumor bearing mice. (**A**) *Left:* Representative FACS plots of total CD4 T cells (CD45^+^CD11b^-^CD11c^-^CD19^-^CD3^+^CD8^-^CD4^+^) and *right:* the corresponding violin plots quantifying the proportion of CD44^+^ PD-1^+^ CD4 T cells. (**B**) *Left:* Representative FACS plots of CD8 T cells (CD45^+^CD11b^-^ CD11c^-^CD19^-^CD3^+^ CD8^+^CD4^-^) and *right:* the corresponding violin plots quantifying the proportion of CD44^+^ PD-1^+^ CD8 T cells. (**C**) *Left:* Representative FACS plots of CD45^+^CD11b^+^ cells. *Center:* histograms showing PD-L1 expression in CD11b^+^ cells. *Right:* quantification of gMFI of PD-L1^+^ CD11b^+^ cells. (**D**) Kaplan Meier survival curve for indicated treatment group. (**A-B**) 2-way ANOVA (*Ern1^f/f^ n* =14, *Ern1^f/f^ Mrp8^Cre^ n* =14). (**C**) 2-way ANOVA (*n* =6/group). (**D**) Log-Rank (Mantel Cox) test, P-value and group numbers provided.

To test this, we induced *Ern1^f/f^* and *Ern1^f/f^ Mrp8^Cre^* mice with autochthonous HGSOC and treated them bi-weekly with i.p. injections of anti-PD-1 or its corresponding IgGa isotype control antibody for four weeks. Consistent with multiple reports in HGSOC patients (Hamanishi et al., 2015), PD-1 blockade had no impact on the overall survival of *Ern1^f/f^* mice bearing these aggressive tumors. However, *Ern1^f/f^ Mrp8^Cre^* mice receiving anti-PD-1 therapy demonstrated a significant increase in overall survival compared with mice of the same genotype receiving control antibody, and with either treatment group in *Ern1^f/f^* hosts (**Fig 5D**). Strikingly, 50% of the *Ern1^f/f^ Mrp8^Cre^* mice that received anti-PD-1 therapy survived more than 150 days post-tumor initiation, suggesting successful long-term control of HGSOC. To confirm this, we harvested spleens from surviving mice for antigen-recall assays using lysates from primary autochthonous tumors, and reactivity was measured by IFN-γ ELiSPOT. Splenocytes from long-term survivors were significantly more responsive when exposed to HGSOC tumor lysates than to control lysates or vehicle (**Fig S5C**). These data indicate that anti-PD-1 therapy combined with restriction of IRE1α signaling in neutrophils unleashes early T cell-mediated immune control of HGSOC and the development of durable protection that prevents disease recurrence.

## Discussion

Here we uncover that the ER stress sensor IRE1α endows neutrophils with marked protumorigenic function in an autochthonous model of HGSOC. Our analysis provides a novel insight to the role of neutrophils in HGSOC tumorigenesis, progression, and immune escape. We have demonstrated that neutrophils rapidly infiltrate the primary tumor while undergoing activation of the IRE1α-XBP1s arm of the UPR. We also determined that deleting IRE1α in neutrophils delays the onset of primary tumors and the progression of metastatic disease by eliciting adaptive T cell responses that can be effectively enhanced upon PD-1 blockade immunotherapy.

Cancer cells in emerging tumor lesions require external support for early proliferation and angiogenic switch, then in later stages, immune escape and metastasis. As early infiltrators with robust tissue regenerative capabilities, neutrophils are key in fulfilling those pro-tumor roles. While additional research is warranted to define how IRE1α may orchestrate these functions in TANs, our study reveals that by genetic knockout of this ER stress sensor, the fundamental events of tumorigenesis are delayed and T cell function against HGSOC is improved. Indeed, the protective effect of IRE1α deletion in neutrophils is completely dependent on anti-tumor T cell activity that is absent in wild type mice. Our findings also suggest that targeting neutrophils in the earliest stages of tumor progression is key to invigorating antigen-dependent T cell responses in HGSOC, a strategy that would be insufficient in immune-escaped metastatic disease. We noted substantial immune reactivity when lymphocytes from HGSOC-bearing mice were exposed to autochthonous primary tumor lysates. However, these antigens appear to be absent in a cell line derived from late-stage metastatic HGSOC.

Abundant neutrophil infiltration into tumor tissues is associated with poor prognosis and reduced therapeutic response in various cancers (Jaillon et al., 2020). Presently, there are early phase clinical trials attempting to impede neutrophil recruitment to the tumor by chemokine blockade as well as functional inhibition of tumor-associated myeloid cells (Goldstein et al., 2021; Melisi et al., 2021). Furthermore, the selective IRE1α pharmacological inhibitor ORIN1001 is currently being tested in patients with different solid cancers (Gabrail, 2021; Li et al., 2024). The data presented in this study indicate that those interventions may be optimized in combination with other immunotherapies for improved HGSOC treatment. Crucially, in autochthonous HGSOC, anti-tumor T cell activity is virtually non-existent when IRE1α expression in neutrophils is intact. Furthermore, we demonstrated that reprogramming the myeloid compartment via IRE1α deletion in neutrophils is critical for the optimal responses to anti-PD-1 therapy. These data indicate that targeting both the myeloid and T cell compartment is fundamental for the effective immunotherapeutic management of HGSOC.

The mechanisms by which ER stress-programmed TANs influence anti-tumor T cell responses in primary autochthonous ovarian tumors are yet to be determined. Our data indicate that the influence of neutrophils on T cells occurs at the earliest phases of tumorigenesis, and due to extremely limited tissue volume and cellularity at the neoplastic stage, we were unable to recover sufficient immune infiltrates to analyze their phenotypes or global molecular profiles. Once primary tumors are firmly established, we may only be observing the vestiges of lymphocyte engagement as evidenced in Figure S4. Analysis of intratumoral neutrophils is further complicated by the heterogenous nature of the autochthonous HGSOC model utilized, where disease initiation and progression are highly variable due to disparities in oncogene incorporation after electroporation. While this phenotypic diversity is useful in replicating clinical realities of cancer, it challenges the analysis of phenotypic changes induced in intratumoral neutrophils lacking IRE1α.

Taken together, our study reveals that IRE1α ablation in neutrophils stimulates early adaptive immune responses that restrain the initiation and progression of primary HGSOC. We show that ablation of IRE1α in neutrophils, combined with anti-PD-1 therapy, elicits robust anti-tumor immunity, increased survival, and durable protection against disease relapse in 50% of treated hosts. These findings provide novel molecular insights for future immunotherapeutic strategies that may improve the clinical management of HGSOC.

## ACKNOWLEDGMENTS

We are grateful to all members of the Cubillos-Ruiz laboratory, former and present, for their valuable suggestions and critical review of this manuscript. We thank the Flow Cytometry Core Facility at Weill Cornell Medicine for their excellent assistance with multiple analyses. We also thank the Scott Lowe Laboratory for generously sharing the MP cell line. This work supported by NIH grants R01 NS114653, R01 CA271619, and R01 CA282072 (J.R.C-R.); U.S. Department of Defense Ovarian Cancer Research Program grants W81XWH2010191, W81XWH-16-1-0438, W81XWH-22-OCRP-IIRA, W81XWH2110478, and W81XWH2110357 (J.R.C-R.); NIH F31CA257631 (A.E.); The AACR-Bristol Myers Squibb Immuno-Oncology Research Fellowship (S.-M.H.); The Cancer Research Institute-Irvington Institute Postdoctoral Fellowship (C-S.C, C.S.); and the Ovarian Cancer Research Alliance (C.S).

## Methods

### Mouse model of autochthonous HGSOC

Female mice were anesthetized with isoflurane and kept at 37°C by a water recirculation platform. The surgical site was scrubbed with a povidone-iodine scrub and washed with 70% ethanol. A paramedian abdominal incision was made to access the left ovary and 25 μl of the plasmid mixture were injected into the ovarian bursa with a 30-gauge needle. The liquid-containing ovary was clamped tightly with a tweezer electrode and two poring pulses were given (50V, 30-ms length, 450-ms intervals) followed by 5 transfer pulses (60V, 50-ms length, 450-ms intervals) using an *in vivo* electroporator (NEPAGENE NEPA21 type II electroporator). Following electroporation of the plasmid mixture into the ovary, the peritoneal cavity was sutured, and the skin was closed with staples. The mice were maintained at 37°C until waking. Pain management was completed with meloxicam injections for 3 days following surgery. The genetic elements used in this model include a sleeping beauty transposable *MYC* overexpression vector and a vector co-expressing Cas9 and single-guide RNA targeting *TP53*, as previously described (Paffenholz et al., 2022). Host survival experiments were monitored over time and defined as ‘post-electroporation’ in all cases. Tumor formation and growth was assessed by abdominal palpation, and size was determined by caliper measurements. HGSOC survival endpoints were defined as when a tumor reached 15x15 mm by caliper measure, and by animal welfare assessment. Mouse welfare was carefully monitored to detect changes in body condition (e.g. fur appearance or abdominal distension), and animals in distress and/or lethargic condition were considered at the humane endpoint. Mice reaching either of these two endpoints were humanely euthanized.

### Cell Lines

ID8-*Defb29/Vegfa* and PPNM (*Trp53^-/-R172H^Pten^-/-^Nf1^-/-^Myc^OE^*) cells were cultured and used as previously described (Conejo-Garcia et al., 2004a; Iyer et al., 2021). These cell lines were obtained from Dr. J. Conejo-Garcia and Dr. R Weinberg respectively, under MTA. Cultured ID8-*Defb29/Vegfa* cells were implanted in mice i.p. by injecting 1.5 x 10^6^ cells suspended in 200 μl of sterile PBS. Metastatic progression and ascites accumulation were measured over time by weighing mice weekly. MP cells were cultured in Dulbecco’s modified Eagle’s medium supplemented with 10% FBS and 100 IU/ml penicillin/streptomycin at 37°C with 5% CO_2_. 1 x 10^6^MP cells were suspended in 200 μl of sterile PBS and injected i.p. into mice. Alternatively, 1 x 10^6^ PPNM cells were suspended in 200 μl of a 1:1 solution of PBS and Matrigel (Corning, Cat# 47743-716) and injected i.p. into mice. Tumor burden and metastatic progression of MP tumor bearing mice were evaluated by bioluminescent imaging. Tumor-bearing mice were i.p. injected with VivoGlo luciferin (2 mg/mouse. Promega, Cat# P1042) and then imaged on the Xenogen IVIS Spectrum In Vivo system housed at the Weill Cornell Research Animal Resource Center. In all cases, host survival was monitored over time. Cell lines were supplemented with prophylactic plasmocin (Invivogen, Cat# ant-mpt) and routinely assessed for mycoplasma contamination.

### Mice

Female mice were housed in pathogen-free microisolator cages at the animal facilities of Weill Cornell Medicine and used at 8-12 weeks of age. C57BL/6J, B6.Cg-Tg (S100A8-cre,-EGFP)1Ilw/J (Mrp8^Cre^), and B6.129P2-Lyz2^tm1(cre)Ifo^/J (LysM^Cre^) were acquired from The Jackson Laboratory. Conditional knockout mice were generated by breeding *Ern1^f/f^* mice with the *Mrp8^Cre^* or *LysM^Cre^*, for selective deletion in neutrophils and in macrophages respectively(Abram et al., 2014; de Boer et al., 2003). All mice were on the C57BL/6J background. Mice were handled in compliance with institutional animal care and use committee guidelines under protocol 2011-0098.

### Immunophenotyping and primary neutrophil isolation

Single-cell suspensions were generated from bone marrow, spleen, peripheral blood, ascites, omentum, and primary ovarian tumors. Bone marrow cells were isolated by flushing the femur and tibia with 10ml of complete RPMI (RPMI1640 supplemented with 10% FBS, L-glutamine, sodium pyruvate, HEPES, nonessential amino acids, β-mercaptoethanol, and penicillin/streptomycin). Peripheral blood cells were isolated by cardiac puncture. Peritoneal ascites were isolated by peritoneal lavage with 5ml of PBS. Spleen, omentum, and ovarian tumor cells were isolated by incubating the tissues with Collagenase D (2 mg/ml) and DNase I (10 ug/ml) at 37°C for 30 mins and then pushed through a 70um cell strainer and washed with RPMI +10% FBS to create a single-cell suspension. Red blood cell removal was performed on all tissue samples by incubating single cell suspensions for 3 minutes with ACK lysis buffer, then washing with 10ml of ice-cold PBS. Lymphocytes sorted from solid tumor samples were further purified by percoll isolation as described in Swamydas *et al (Swamydas and Lionakis, 2013)*. Cells were then resuspended in FACS buffer (PBS with 0.5% BSA and 2mM EDTA) and FC-γ receptor blocked with TruStain FcX (anti-mouse CD16/32), followed by surface markers were stained at 4°C for 30 minutes in the dark. Cells were then washed and resuspended in PBS and stained with DAPI for live/dead cell identification. For intracellular staining, single cell suspensions were stained with violet Live/dead stain for 4°C for 30 minuntes in the dark, following surface marker staining. Cells were then fixed using the Foxp3 intracellular staining kit (eBiosciences, Cat: 00-5523-00) followed by intracellular staining. For cytokine expression assays, single cell suspensions generated from primary ovarian tumors were stimulated *ex vivo* with phorbol 12-myristate 13-acetate (PMA), ionomycin, and brefeldin A for 5 hours before intracellular staining with the Foxp3 protocol. Flow cytometry was performed on a Foressa-X20 (BD Biosciences), and data analysis was completed with FlowJo software (TreeStar). FACS sorting was performed in the Weill Cornell Medicine Flow Cytometry Core Facility and Neutrophils (CD45^+^CD11c^-^CD19^-^F4/80^-^CD11b^+^Ly6G^Hi^Ly6C^Lo^) were sorted on a BD FACSAria-II or FACSymphony A5 and processed for RNA isolation. Flow cytometry antibodies used in this study are described in Supplemental Table 1

### Nucleic acid isolation and quantitative RT-PCR analyses

Mouse total RNA was extracted using the RNeasy plus mini kit (Qiagen, Cat# 74134) or the Arcturus PicoPure RNA isolation kit (Applied Biosystems, Cat# KIT0204), according to manufacturer’s instructions. 0.1-0.25 μg of purified RNA was reverse transcribed to generate cDNA using the qScript cDNA synthesis kit (Quantabio, Cat# 95047). Quantitative RT-PCR was performed with PerfeCTa SYBR green fast-mix (Quantabio, Cat# 95071) and run using the QuantStudio 6 Flex real-time PCR system (Applied Biosciences). Gene expression values were calculated using the comparative threshold cycle method and normalized to *Actb* as endogenous controls. Primers used in this study are described in Supplemental Table 2.

### In vivo treatments

*Neutrophil depletion.* The ovaries of wild type C57BL/J6 mice were electroporated as described in Figure 1A. After 10 days, mice were i.p. treated at 200 μg/mouse of an anti-Ly6G (Clone: 1A8, BioXCell, Cat: BP0075-1) or an IgG2a isotype control antibody (Clone: 2A3, BioXCell, Cat: BP0089). Mice were treated every 48 hours for five weeks. Antibodies were prepared as described by the manufacturer’s instructions.

*T cell depletion*. The ovaries of *Ern1^f/f^ Mrp8^Cre^* or *Ern1^f/f^* mice were electroporated, and after 10 days, mice were i.p. treated with 150 μg/mouse each of an anti-CD4 (Clone: GK1.5, BioXCell, Cat: BE0003-1) and anti-CD8 (Clone: HB-129, BioXCell, Cat: BE0118) or an IgG2b isotype control (Clone: LFT-2, BioXCell, Cat: BE0090). Mice were treated every 3 days until endpoint.

*Checkpoint therapy*: The ovaries of *Ern1^f/f^ Mrp8^Cre^* or *Ern1^f/f^* mice were electroporated, and after 14 days, mice were i.p. treated at 200 μg/mouse of an anti-PD1 (clone: RMP1-14, BioXcell, Cat: BE0146) or IgG2a isotype control antibody (Clone 2A3, BioXCell, Cat: BE0089). Mice were treated bi-weekly for four weeks.

### ELISpot Assays

BMDCs were differentiated by incubation of total bone marrow cells in cRPMI with recombinant GM-CSF (20 ng/ml). After 6 days, culture medium was replenished then non-adherent cells were harvested on day 7. MP cells were collected as described under *Cell Lines*. Primary autochthonous tumors were resected and chemically digested with collagenase D (2 mg/ml) and DNase I (10 ng/ml) to produce a single-cell suspension. Protein lysates were generated by rapid freeze/thaw cycles and quantified with a standard curve colorimetric assay (Pierce BCA protein assay kit, ThermoScientific Cat: 23225). BMDC and tumor protein lysates were derived from *Ern1^f/f^* female mice. Primary splenocytes and bone marrow cells were seeded at 0.8 x 10^6^ cells/well and incubated with 200ug/well of the specified lysates for 24 hours. IFNγ ELISpot assays were performed using pre-coated 96-well plates (Immunospot Cat: mIFNg-2M/2) according to the manufacturer’s instructions. Spots were counted with the S6 Universal M2 (Immunospot).

### Statistical Analyses

All Statistical analyses were performed with GraphPad Prism 10. Significance of survival was calculated using the Log-rank test. Comparisons between two groups were made with unpaired two-tailed Student’s *t*-test or Dunnett’s multiple comparisons test. Multiple comparisons were completed with one-way ANOVA. Comparisons of multiple groups were done with Sidak’s multiple comparisons test. The significance of pairwise correlation analysis was calculated with Pearson’s correlation coefficient. All data are presented as mean +/- SEM unless otherwise stated. Exact P-values are shown on all graphs, where a P-value of <0.05 was considered statistically significant.

