## Supplemental Figures and Tables for "High-grade serous ovarian cancer development and anti-PD-1 resistance is driven by IRE1α activity in neutrophils"

SUPPLEMENTARY FIGURE 1

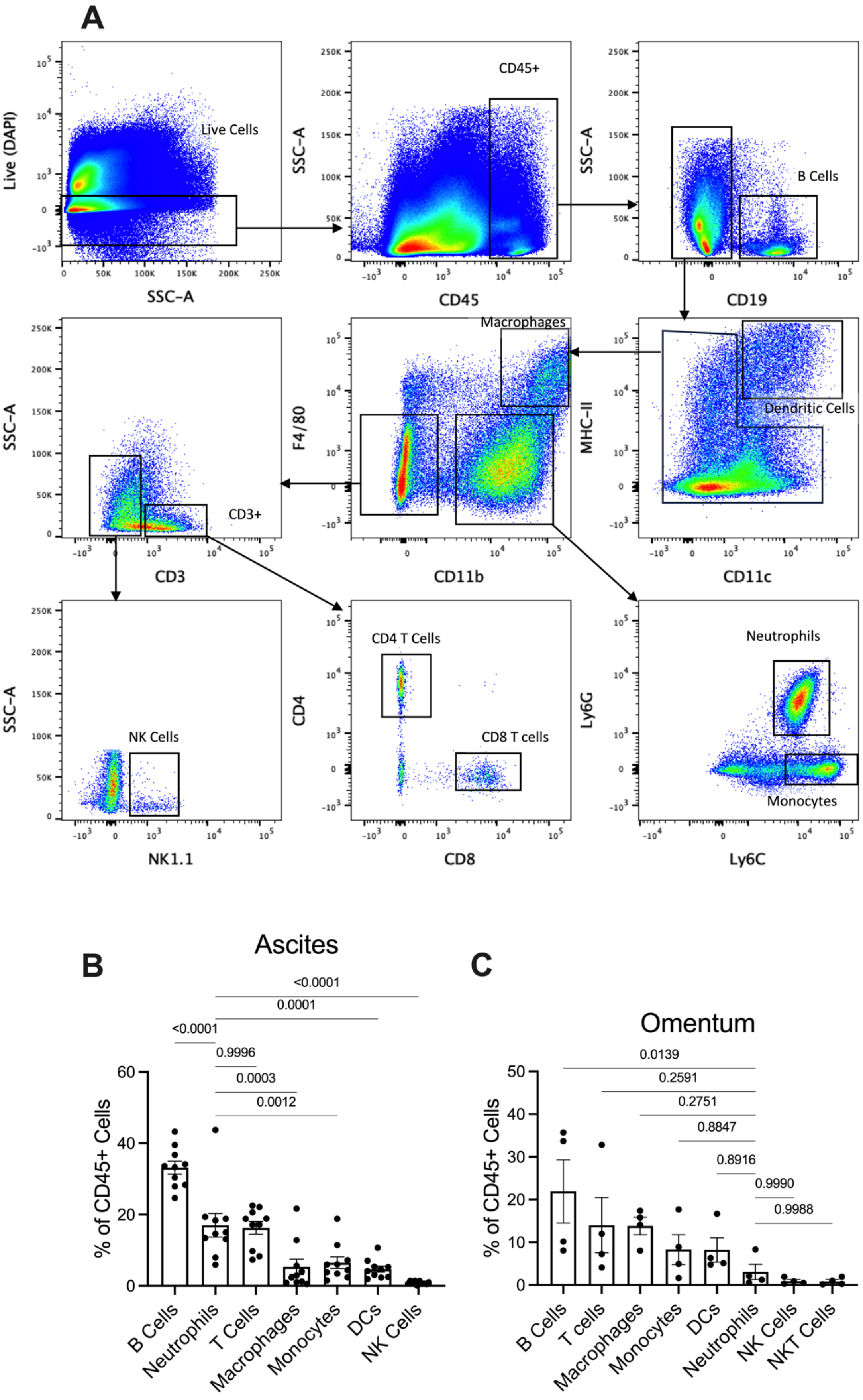

Supplementary Figure 1: C57BL/6J mice were induced with autochthonous HGSOC as described in figure 1A (**A**) Flow cytometry gating strategy to identify immune populations in HGSOC-bearing mice. (**B-C**) Bar graphs of the flow cytometry-based quantification of immune populations in the (**B**) ascites and (**C**) metastasized omenta of HGSOC-bearing mice. Non-hemorrhagic ascites and non-metastasized omenta were excluded from the analysis. (**B**) 1-way ANOVA (*n* =10). (**C**) 1-way ANOVA (*n* =4). P-values provided.

SUPPLEMENTARY FIGURE 2

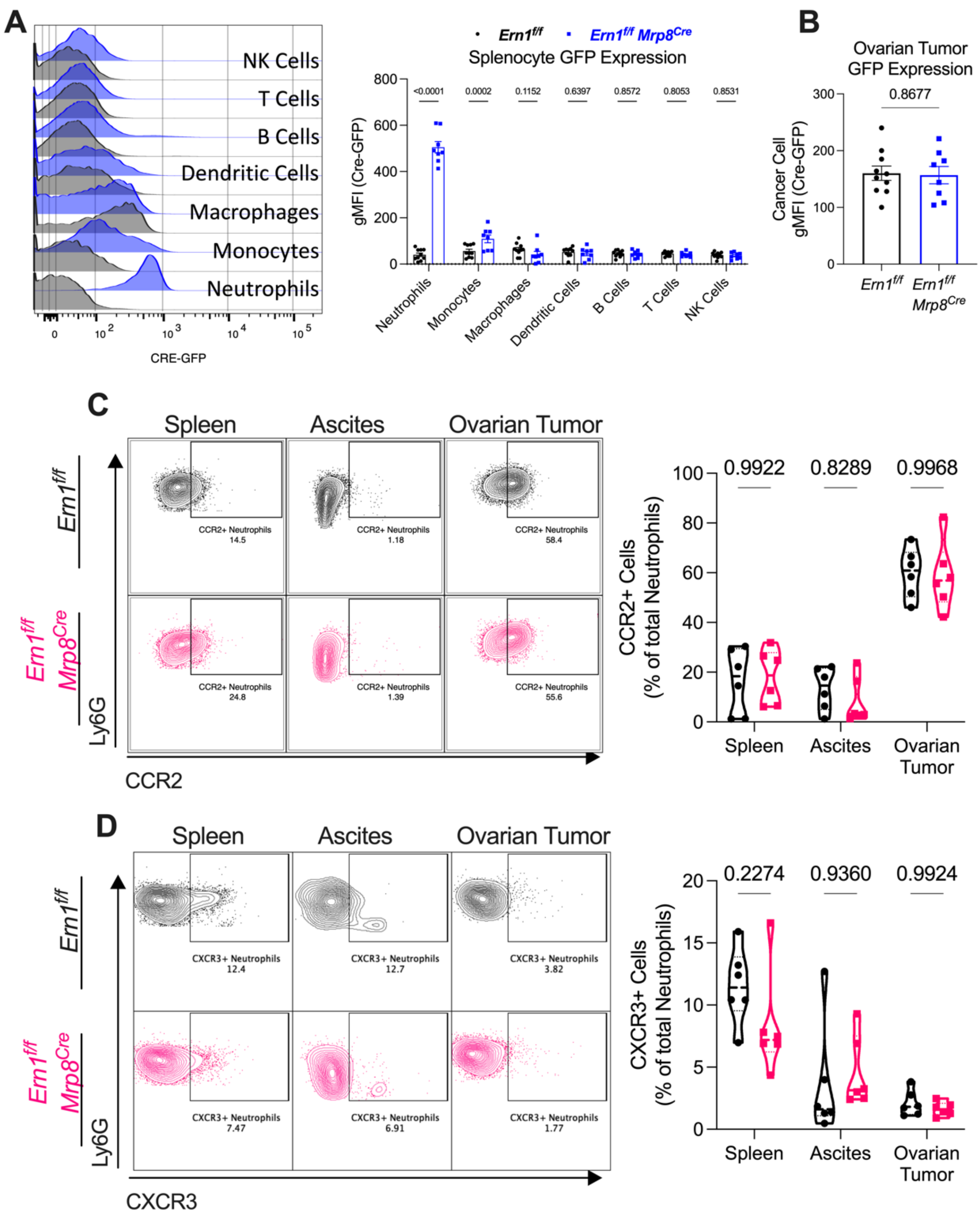

Supplementary Figure 2: Analysis of *Ern1*$^{f/f}$ and *Ern1*$^{f/f}$ *Mrp8*$^{Cre}$ female mice with autochthonous HGSOC. (**A**) *Right:* Bar graph of geometric mean fluorescent intensity (gMFI) of CRE-GFP expression from the indicated populations and *left:* accompanying representative histograms. (**B**) Bar graph of gMFI of CRE-GFP expression from $CD45^{-}SSC^{Hi}$ cells from the primary ovarian tumor of the indicated groups and accompanying histograms. (**C-D**) Violin plots of (**C**) $CCR2^{+}$ and (**D**) $CXCR3^{+}$ neutrophils ($CD45^{+}CD19^{-}CD11c^{-}NK1.1^{-}CD3^{-}CD11b^{+}Ly6G^{Hi}Ly6C^{Lo}$) from tumor-bearing mice and accompanying representative FACS plots. A) 2-way ANOVA (*Ern1*$^{f/f}$ *n* =10, *Ern1*$^{f/f}$ *Mrp8*$^{Cre}$ *n* =8). (**B**) Unpaired t-test (*Ern1*$^{f/f}$ *n* =10, *Ern1*$^{f/f}$*Mrp8*$^{Cre}$ *n* =8). (**C-D**) 2-way ANOVA (*Ern1*$^{f/f}$ *n* =6, *Ern1*$^{f/f}$*Mrp8*$^{Cre}$ *n* =6). P-values provided.

SUPPLEMENTARY FIGURE 3

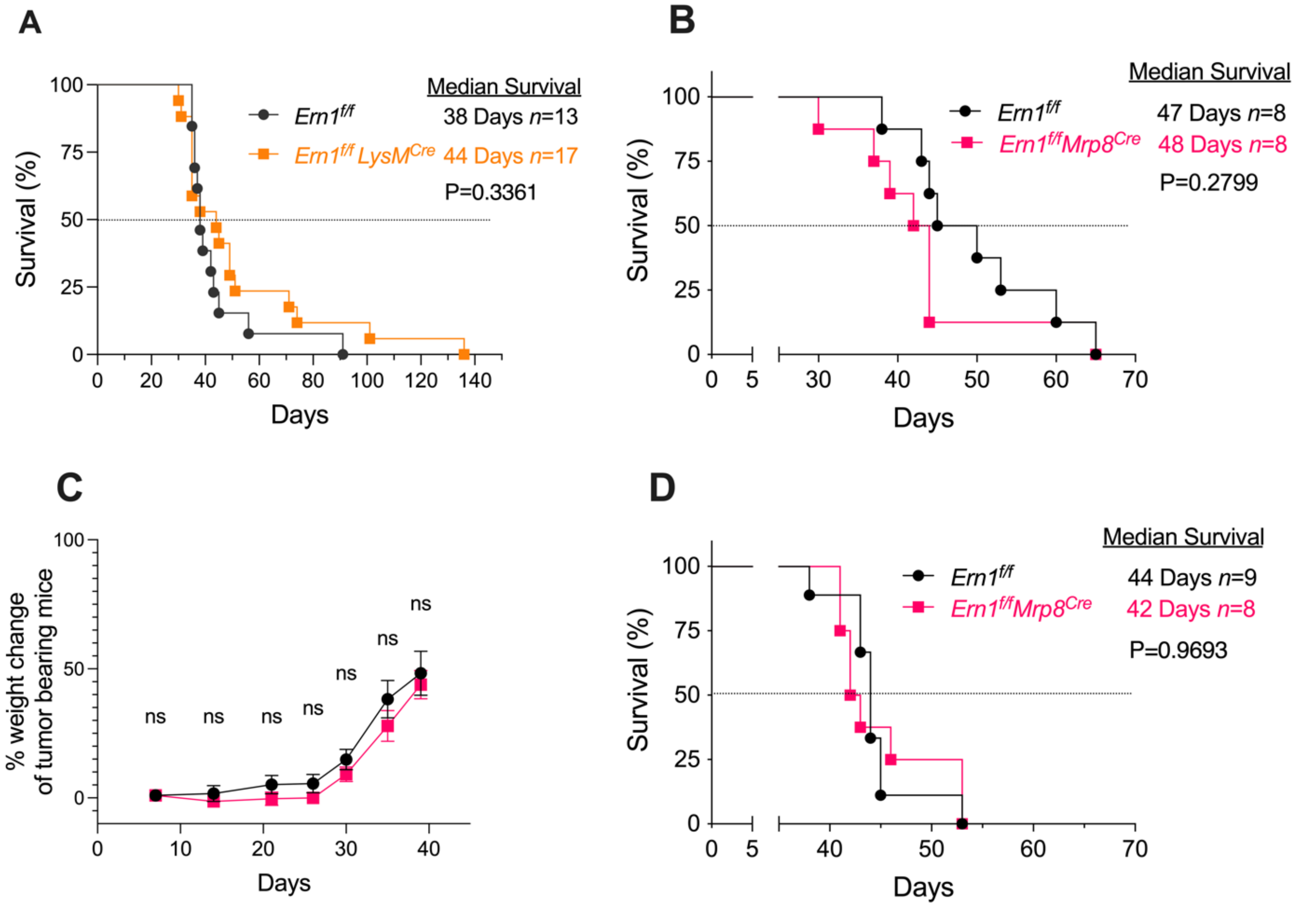

Supplementary Figure 3: (**A**) Kaplan-Meier survival curve of mice bearing autochthonous HGSOC. (**B-D**) Female mice were injected i.p. with syngeneic ovarian cancer as indicated. (**B**) Kaplan-Meier survival curve of mice challenged with the transplantable PPNM ovarian cancer cell line. (**C-D**) Female mice were challenged with the ID8-*Defb29/Vegfa* ovarian tumor cell line: (**C**) ascites accumulation over the course of tumor growth (**D**) Kaplan-Meier survival curve of tumor-bearing mice. (**A-B, D**) Log-Rank (Mantel Cox) test, P-value and group numbers provided. (**C**) Two-way ANOVA ($Ern1^{f/f}$ $n$ =9, $Ern1^{f/f}$ Mrp8$^{Cre}$ $n$ =8).

SUPPLEMENTARY FIGURE 4

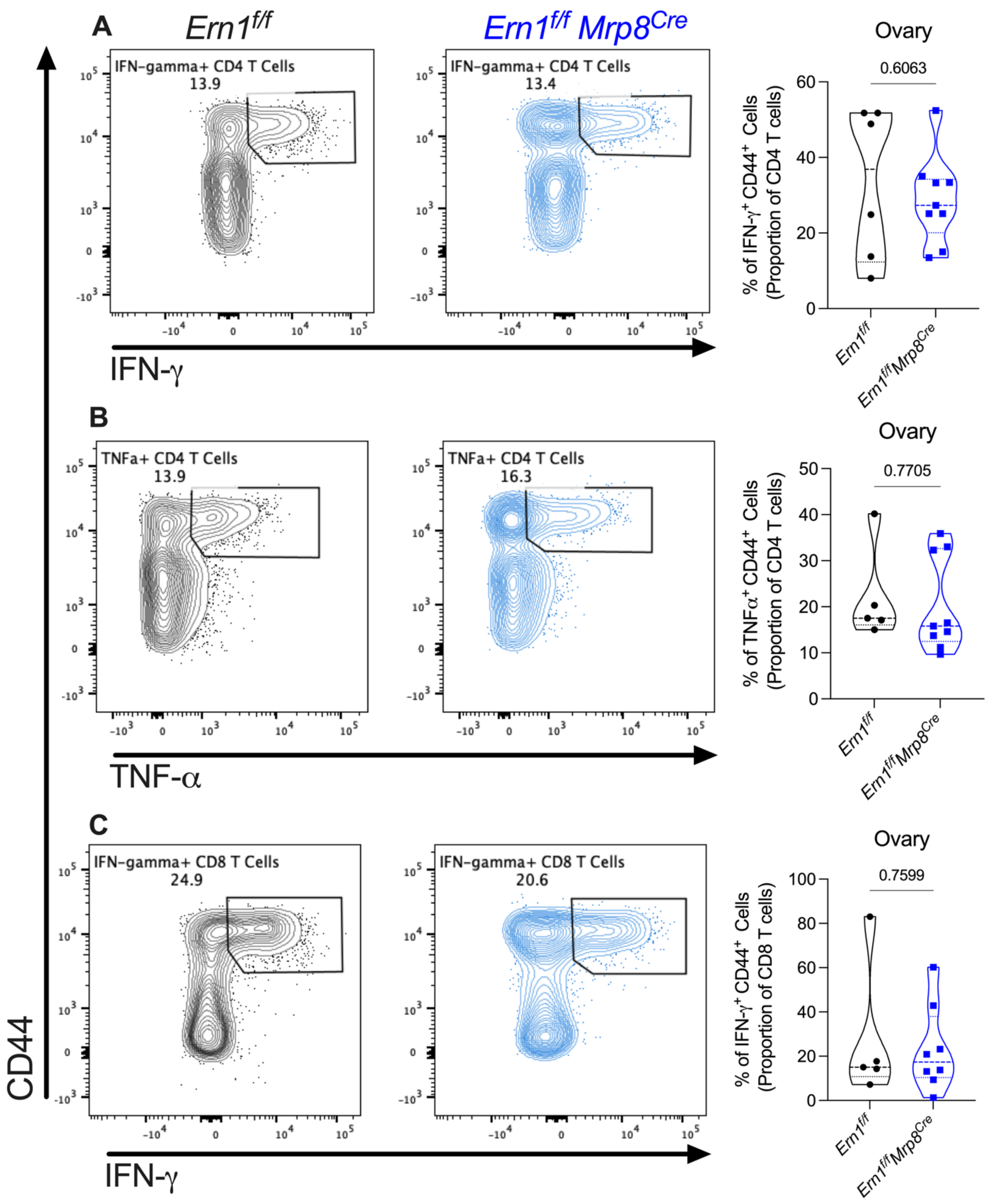

Supplementary Figure 4: (**A-C**) *Left:* Representative FACS plots and *right:* violin plots with the proportion of CD4 or CD8 T cells from primary ovarian tumors expressing IFNγ or TNFα. Unpaired Student's t test (*Ern1$^{f/f}$ n* =6, *Ern1$^{f/f}$Mrp8$^{Cre}$ n* =9). P-values provided.

SUPPLEMENTARY FIGURE 5

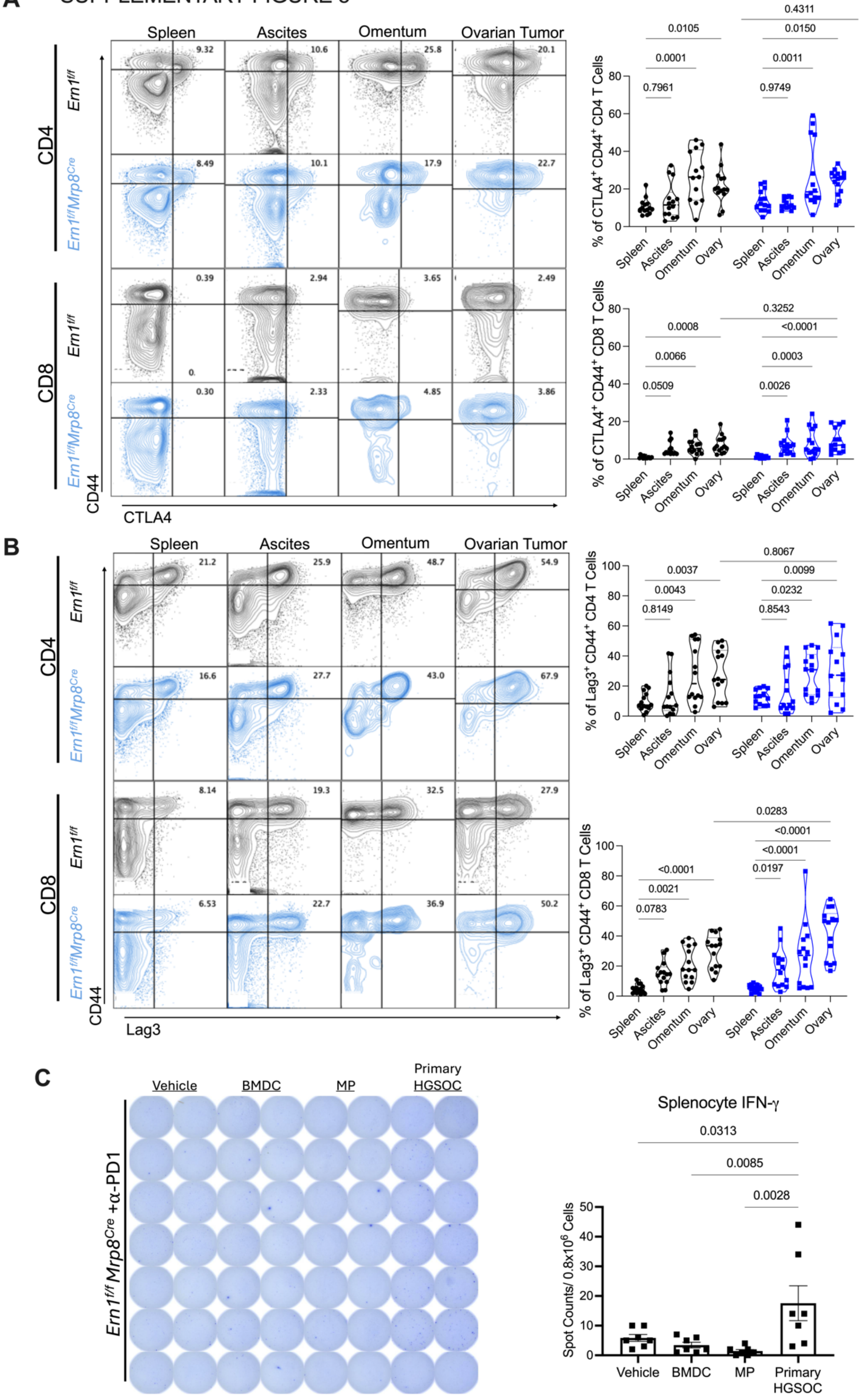

Supplementary Figure 5: (**A-B**) Analysis of T cells ($CD45^{+}CD11b^{-}CD11c^{-}CD19^{-}CD3^{+}$) from *Ern1^f/f^* and *Ern1^f/f^Mrp8^Cre^* female mice developing autochthonous HGSOC. (**A-B**) *Left*: Representative FACS plots and *right:* corresponding violin plots quantifying the proportion of $CTLA4^{+}CD44^{+}$ CD4 and CD8 T cells and (**B**) $Lag3^{+}CD44^{+}$ CD4 and CD8 T cells. (**C**) *Left:* Representative image of IFN-γ ELISpot assay and *right:* Quantification of results. Total splenocytes or bone marrow cells were seeded in a 96-well plate and stimulated with the indicated lysates. Each point represents an average of two plate replicates. (**A-B**) 2-way ANOVA ($Ern1^{f/f}$ *n* =14, $Ern1^{f/f}Mrp8^{Cre}$ *n* =14). (**C**) One-way ANOVA (*n* =7). P-values provided.

Supplementary Table 1

| Fluorophore | Target | Clone | Manufacturer | Catalog Number |
|---|---|---|---|---|
| PE-CF594 | CD45 | 30-F11 | BD Biosciences | 562420 |
| BV605 | CD3 | 17A2 | Biolegend | 100237 |
| APC-Cy7 | CD4 | GK1.5 | Biolegend | 100414 |
| BUV396 | CD8a | 53-6.7 | BD biosciences | 563786 |
| BV421 | CD44 | IM7 | Biolegend | 103040 |
| PE-Cy7 | CD19 | 1D3 | Tonbo Biosciences | 60-0193-U025 |
| PE-Cy7 | CD11c | N418 | Biolegend | 117318 |
| PE-Cy7 | CD11b | M1/70 | Tonbo Biosciences | 60-0112-U100 |
| FITC | GR-1 | RB6-8C5 | Biolegend | 108406 |
| PerCP-Cy5.5 | TNF-$\alpha$ | MP6-XT22 | Biolegend | 506322 |
| BUV737 | IFN-$\gamma$ | XMG1.2 | BD biosciences | 564693 |
| BV711 | PD-1 | 29F.1A12 | Biolegend | 135231 |
| AF700 | Ki67 | 16A8 | Biolegend | 652419 |
| BV786 | Lag3 | C9B7W | Biolegend | 125219 |
| PerCP-Cy5.5 | CD62L | MEL-14 | Biolegend | 104432 |
| APC | CTLA4 | UC10-4F10-11 | Tonbo Biosciences | 20-1522-U100 |
| PerCP Cy5.5 | CD4 | RM4-5 | Tonbo Biosciences | 65-0042 |
| BV711 | CD11c | N418 | Biolegend | 117349 |
| PE-Cy7 | CD44 | IM7 | Biolegend | 103030 |
| PE-Cy7 | F4/80 | BM8 | eBioscience | 25-4801-82 |
| BV786 | Ly6C | HK1.4 | Biolegend | 128041 |
| B510 | Ly6G | 1A8 | Biolegend | 127633 |
| APC | I-A/I-E | M5/114.15.2 | eBioscience | 17-5321-82 |
| BUV737 | NK1.1 | PK136 | BD biosciences | 741715 |
| BV421 | CD11b | M1/70 | Biolegend | 101251 |
| PE Dazzle 594 | CD19 | 6G5 | Biolegend | 115554 |
| Alexa Fluor 700 | CD45 | 30-F11 | Biolegend | 103128 |
| PE-Cy7 | CCR2 | SA203G11 | Biolegend | 150611 |
| BV605 | CXCR3 | 493 | BD bioscience | 745114 |
| BUV 737 | PD-L1 | MIH5 | BD biosciences | 741877 |
| PerCP Cy5.5 | CD11b | M1/70 | Biolegend | 101228 |

Supplementary Table 2

| Species | Gene | Primer Direction | Sequence 5' - 3' | Purpose |
|---|---|---|---|---|
| Mouse | *Actb* | Forward | CTCAGGAGGAGCAATGATCTTGAT | RT-qPCR |
| | | Reverse | TACCACCATGTACCCAGGCA | |
| Mouse | *Xbp1s* | Forward | AAGAACACGCTTGGGAATGG | RT-qPCR |
| | | Reverse | CTGCACCTGCTGCGGAC | |
| Mouse | *Xbp1* | Forward | GACAGAGAGTCAAACTAACGTGG | RT-qPCR |
| | | Reverse | GTCCAGCAGGCAAGAAGGT | |
| Mouse | *Dnajb9* | Forward | TAAAAGCCCTGATGCTGAAGC | RT-qPCR |
| | | Reverse | TCCGACTATTGGCATCCGA | |
| Mouse | *Hspa5* | Forward | TCATCGGACGCACTTGGAA | RT-qPCR |
| | | Reverse | CAACCACCTTGAATGGCAAGA | |
| Mouse | *Ddit3* | Forward | GTCCCTAGCTTGGCTGACAGA | RT-qPCR |
| | | Reverse | TGGAGAGCGAGGGCTTTG | |
